# Direct measurements of luminal Ca^2+^ with endo-lysosomal GFP-aequorin reveal functional IP_3_ receptors

**DOI:** 10.1101/2023.07.11.547422

**Authors:** B Calvo, P Torres-Vidal, A Delrio-Lorenzo, C Rodriguez, FJ Aulestia, J Rojo-Ruiz, BM McVeigh, V Moiseenkova-Bell, DI Yule, J Garcia-Sancho, S Patel, MT Alonso

**Author notes:** These authors contributed equally to this work (B Calvo, P Torres-Vidal).

## Abstract

Endo-lysosomes are considered acidic Ca^2+^ stores but direct measurements of luminal Ca^2+^ within them are limited. Here we report that the Ca^2+^-sensitive luminescent protein aequorin does not reconstitute with its cofactor at highly acidic pH but that a significant fraction of the probe is functional within a mildly acidic compartment when targeted to the endo-lysosomal system. We leveraged this probe (ELGA) to report Ca^2+^ dynamics in this compartment. We show that Ca^2+^ uptake is ATP-dependent and sensitive to blockers of endoplasmic reticulum Ca^2+^ pumps. We find that the Ca^2+^ mobilizing messenger IP_3_ which typically targets the endoplasmic reticulum evokes robust luminal responses in wild type cells, but not in IP_3_ receptor knock-out cells. Responses were comparable to those evoked by activation of the endo-lysosomal ion channel TRPML1. Stimulation with IP_3_-forming agonists also mobilized the store in intact cells. Super-resolution microscopy analysis confirmed the presence of IP_3_ receptors within the endo-lysosomal system, both in live and fixed cells. Our data reveal a physiologically-relevant, IP_3_-sensitive store of Ca^2+^ within the endo-lysosomal system.

## INTRODUCTION

The endo-lysosomal (EL) system has emerged as a relevant player in subcellular Ca^2+^ signalling alongside the better characterised endoplasmic reticulum (ER) (Morgan et al., 2011; Patel and Docampo, 2010; Yang et al., 2019). EL Ca^2+^ has been implicated in most of the organelle’s functions, such as late endosome-lysosome fusion, lysosomal exocytosis, and membrane repair. Impaired Ca^2+^ signalling can cause defects in EL trafficking and eventually can provoke lysosome storage diseases (LSDs). These so called acidic Ca^2+^ stores are thought to accumulate high amounts of Ca^2+^ (in the range of 0.4-0.6 mM), similar to the ER (Christensen et al., 2002; Lloyd-Evans et al., 2008). Akin to ER-localised receptor/Ca^2+^ channels for the second messenger IP_3_ which mediate Ca^2+^ release in response to IP_3_-forming hormones, various Ca^2+^-permeable channels have been shown to localize to membranes of the EL system. These include Mucolipin transient receptor potential (TRPML) channels, two pore channels (TPCs) and P_2_X_4_ channels. TRPML1 is an ubiquitously expressed channel specifically activated by phosphatidylinositol 3,5-bisphosphate (PI(3,5)P_2_), a low abundant phosphoinositide enriched in endomembranes, and which is mutated in type IV mucolipidosis (Dong et al., 2010).

The ER accumulates Ca^2+^ via SERCA. In contrast, the mechanism responsible for EL Ca^2+^ uptake has not been yet fully elucidated, mainly because of the technical difficulties associated to measure intraluminal Ca^2+^ in an acidic compartment. It is assumed that the vacuolar-type ATPase (V-ATPase) which is present in the EL membrane and maintains the proton gradient, drives Ca^2+^ into the lysosomal lumen via a type of transporter of H^+^ for Ca^2+^ similar to the Ca^2+^/H^+^ exchanger present in yeast and in some animals (Melchionda et al., 2016; Mindell, 2012). This view has been recently challenged by the proposal that the ER, not the pH gradient, through IP_3_R tunnels Ca^2+^ into the endo-lysosomes (Atakpa et al., 2019; Garrity et al., 2016). But these conclusions were drawn from indirect monitoring of cytosolic Ca^2+^ using conventional organic Ca^2+^ dyes or genetically-encoded Ca^2+^ indicators targeted to the cytosolic site of the EL system (Kilpatrick et al., 2016; Shen et al., 2012) thereby providing only qualitative insights into EL Ca^2+^ uptake.

Direct measurements of luminal Ca^2+^ within the EL system have been challenging due to the highly acidic pH which interferes with Ca^2+^ binding and thus requires careful calibration. In this study, we report the use of the Ca^2+^-sensitive luminescent protein aequorin targeted to the endo-lysosome. In general, aequorin is considered less sensitive to acidic pH than fluorescent genetically encoded Ca^2+^ indicators (Johnson and Shimomura, 1978; Shimomura et al., 1974). But surprisingly, we find that aequorin does not reconstitute well at extreme acidic pH and therefore reports Ca^2+^ in a mildly acidic compartment of the EL system. We show that this compartment accumulates Ca^2+^ via active uptake and releases it via endogenous IP_3_ receptors. Our data extend the actions of IP_3_ to the endo-lysosomal system.

## RESULTS

### ELGA as a novel endo-lysosomal Ca^2+^ indicator

We recently introduced genetically encoded Ca^2+^ indicators based on aequorin fused to GFP (Rodriguez-Garcia et al., 2014; Rodriguez-Prados et al., 2015). Here, we targeted them to the EL system by fusing the N-terminal of GFP-aequorin (GA) to the C-terminal of the tetanus-insensitive vesicle-associated membrane protein (VAMP7), a marker of the late endosomes and secretory lysosome (Pryor et al., 2004; Vats and Galli, 2022) (***Figure 1A***). VAMP-7 is composed of a cytosolic region, an EL transmembrane domain and a luminal C-terminal domain. Therefore, the Ca^2+^ indicators are expected to be confined to the EL lumen (***Figure 1B***). Due to the expected high endo-lysosomal luminal Ca^2+^, we introduced the D119A mutation in aequorin to lower the affinity for Ca^2+^. This probe was dubbed ELGA (for <u>E</u>ndo-<u>L</u>ysosomal <u>G</u>FP-<u>A</u>equorin). We generated a second variant (ELGA1) with additional substitutions (D117A, and D163A) to further reduce the affinity, lowering rate of aequorin consumption and thereby allowing longer recordings (up to ∼1h). Note that GA1 was a variant previously published as GAP1 (Rodriguez-Garcia et al., 2014; Rodriguez-Prados et al., 2015) but renamed here to maintain a comprehensive nomenclature. Both probes can measure Ca^2+^ in the range of 100-1000 µM (***Figure 1 – figure supplement 1A***). The Ca^2+^ sensors were expressed (either transiently or stably) in mammalian cells using a relatively weak herpes simplex virus type 1 (HSV-1) promoter (Geller, 1991), to limit the amount of expressed protein and, hence, to minimize mistargeting.

**Figure 1.**
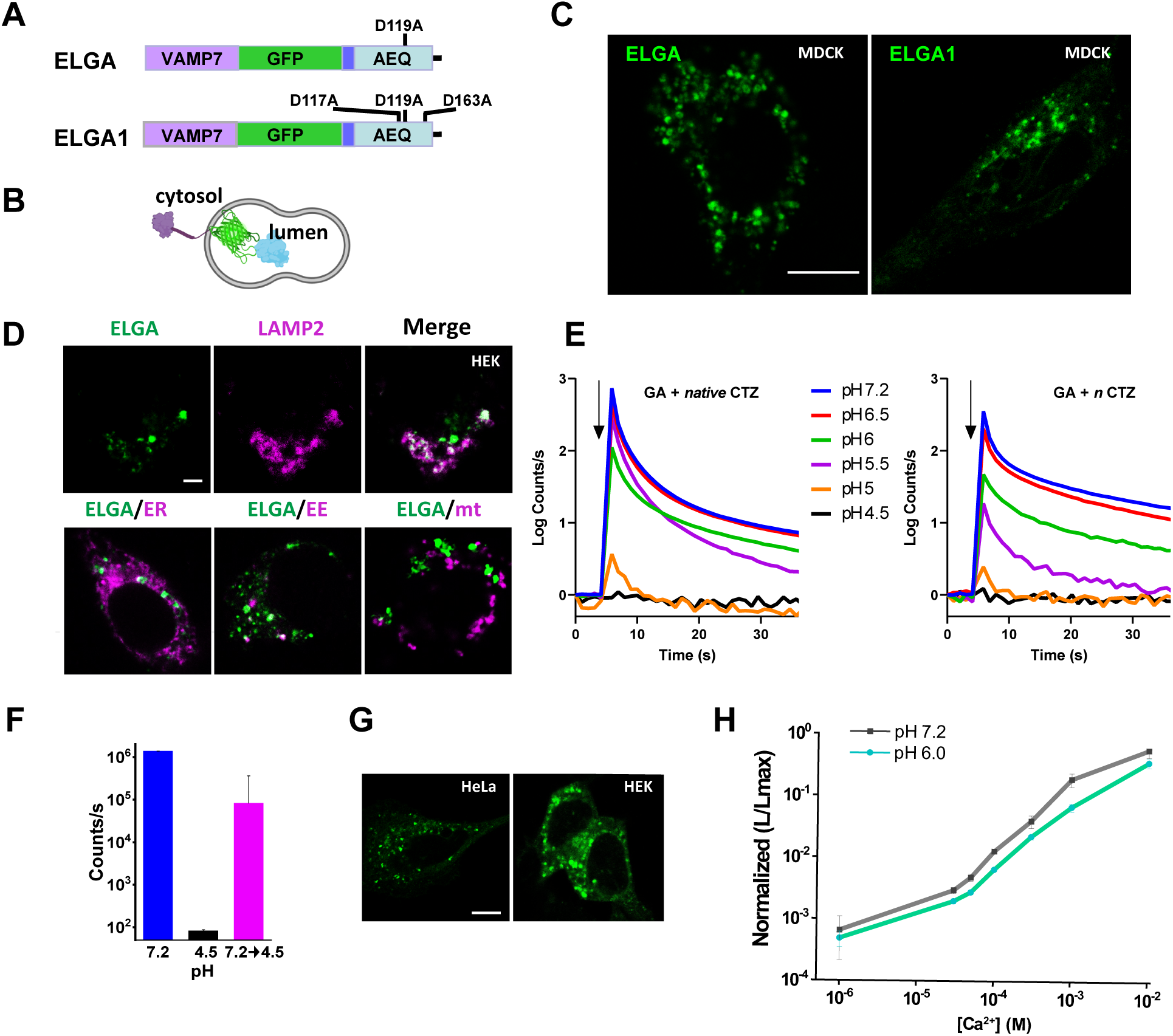
Design, characterization and calibration of endo-lysosomal Ca^2+^ indicators. **(A)** Domain structure and mutations of the endo-lysosomal indicators drawn approximately at scale. VAMP7, vesicle-associated membrane protein 7; AEQ, aequorin; ELGA, endo-lysosomal GFP-Aequorin; ELGA1, endo-lysosomal GAP1. **(B)** Predicted topology of ELGA and ELGA1 Ca^2+^ probes within the endo-lysosome. **(C)** Representative confocal fluorescent images of equatorial plane of fixed MDCK cells transiently transfected with ELGA or ELGA1. Scale bar for both images, 10 µm. **(D) *Upper row:*** ELGA expressing HEK293T cells (*Left*, green) were immunostained with a specific anti-LAMP2 antibody (*Center*, magenta). Green and magenta images are merged (*Right, merge*). ***Lower row:*** Colocalization of ELGA expressing HEK293T cells with the following organellar markers (magenta): ER, er-Ruby-GCaMP210 (ER); early endosome, immunostained with the anti-early endosome antigen 1 (EE); mitochondria, immunostained with anti-TOM20 antibody (mt). Scale bar for all images, 5 µm. **(E)** Effect of acidic pH on GA luminescence emission. *E.coli* recombinant GA apoprotein was reconstituted with native (*left*) or *n* (*right*) coelenterazine (CTZ) for 20 min at the indicated pH, and then subjected to luminescence emission assay triggered with 10 mM CaCl_2_ (arrow) and measured at the same pH. Buffers used were: acetate for range 4.5-5.5 pH; Na-MOPS for 6 and 6.5 pH; and Na-Hepes for 7.2 pH. Each trace is the value normalized with respect to its background emission before addition of Ca^2+^. Each trace is the average of 3-4 independent experiments. **(F)** Luminescence activity of GA1 after regeneration with *n* coelenterazine at 7.2 or 4.5 pH. Peak value (mean ± S.E.M; n=3) obtained from an experiment similar to (E) performed at pHs 7.2 *(blue bar)* and 4.5 *(black bar)* or reconstituted with coelenterazine at pH 7.2 and switched to pH 4.5 to measure luminescence (*pink bar*). **(G)** Expression of ELGA in live HeLa and HEK293T cells. Scale bar, 10 µm. **(H)** Ca^2+^ titration curves of GA1 at pH 6.0 and 7.2. Each value is the mean ± S.E.M of 3-6 measurements obtained from two independent protein preparations.

As shown in ***Figure 1C-supplement movie 1***, ELGA and ELGA1 both localized to punctate structures. To characterize the subcellular distribution of the ELGAs, we performed colocalization analysis with various endogenous organellar markers. ***Figure 1D*** shows representative confocal images of HEK293T cells transfected with ELGA and fixed one day later. The endogenous lysosome-associated membrane protein 2 (LAMP-2) was assayed by immunodetection with a specific antibody (***Figure 1D***). There was colocalization of ELGA with LAMP2, whereas there was no significant overlap with markers for other organelles such as ER (Farrell et al., 2024), early endosome, or mitochondria (***Figure 1D-figure supplement 2 and table supplement 1***). Altogether, these results indicate that the majority of the ELGA indicator is localized to a subset of endo-lysosomes.

### ELGA reports Ca^2+^ in mildly acidic compartments

To analyse the ability of aequorin to report Ca^2+^ at acidic pHs, we compared the maximal luminescence signal triggered by Ca^2+^ in the pH range from 4.5 up to 7.2. Recombinant GA regenerated with coelenterazine showed maximal luminescence activity at pH 7.2 that decreased proportionally at lower pHs (6.5 and 6.0 pH) up to a threshold of 5.5 pH, below which the Ca^2+^ sensor was not functional (***Figure 1E).*** Similar results were obtained with GA1 or with the original aequorin, either reconstituted with native coelenterazine or with its lower affinity *n* analogue (***Figure supplement 3***), indicating that the effect of acidic pH on the luminescent function is independent on the affinity of aequorin for Ca^2+^. This was surprising, given previous calibration/performance studies at acidic pH (Mitchell et al., 2001; Ronco et al., 2015; SantoDomingo et al., 2010). If the aequorin-based Ca^2+^ sensor was first reconstituted at near-neutral pH (7.2) and then, shifted to pH 4.5 for the luminescence assay, the luminescence activity was recovered, suggesting that the lack of light emission is due to the binding of the cofactor to apoaequorin (***Figure 1F)***.

The above findings precluded measurement of Ca^2+^ at highly acidic pH such as that found in lysosomes. But because the pH of endocytic organelles is diverse (e.g., early endosome, pH 6.5; late endosomes, pH 6.0; lysosome, pH 4.5), we reasoned that ELGAs could be used to report Ca^2+^ in less acidic compartments within the EL system. To determine whether ELGAs reside in such compartments, we leveraged the pH sensitivity of the GFP moiety in ELGA. GFP fluorescence is quenched at acidic pH (pKa ∼6). Thus, any detectable GFP fluorescence in live cells should reflect targeting to a compartment with a pH>5 within the range where aequorin can be reconstituted. As shown in ***Figure 1G***, punctate GFP fluorescence was readily detectable in live HeLa and HEK293 cells, both transfected with ELGA.

To assess the properties of the probes at less acidic pH, we performed Ca^2+^ calibrations at pH 7.2 and 6. Essentially similar values were obtained at both pH values, indicating no major effect within this range. As shown in ***Figure 1H-supplement 1A***, the affinity of GA1 for Ca^2+^ was ∼350 µM at pH 7.2.

Taken together, these results identify ELGAs as reporters of Ca^2+^ within mildly acidic compartments.

### Uptake of Ca^2+^ is ATP-dependent and sensitive to blockers of SERCA

In order to directly monitor uptake and release of Ca^2+^ within the EL compartment, we used digitonin-permeabilized HEK293T cells expressing ELGA1 perfused with a cytosol-like solution. As we expected the EL system to be a high Ca^2+^ content store, it needed to be depleted of Ca^2+^ before aequorin was regenerated with its cofactor to avoid its premature consumption. But the mechanisms of store filling are unclear. We first tested Bafilomycin A1 prior to adding the coelenterazine: this compound inhibits the V-type ATPase and has been used widely to disrupt the activity of putative Ca^2+^-H^+^ exchangers. Bafilomycin A1 treatment did not produce a measurable luminescent signal (data not shown), indicating that aequorin had been consumed likely because the target store was not emptied of Ca^2+^. These data are consistent with functional targeting of ELGA to a non-highly acidic compartment.

By contrast, incubation with the reversible SERCA inhibitor tert-butylhydroquinone (tBHQ), produced a robust luminescence with a good signal to noise ratio (***Figure 2A***). Under these conditions perfusion with cytosol-like medium (free [Ca^2+^] = 200 nM) initiated Ca^2+^ uptake at a rate of 8 ± 0.6 µM/s (n=36), reaching a value of [Ca^2+^]_EL_ at the steady-state of 365 ± 21 µM (n=54) in 3 ± 0.6 min (n=7) (***Figure 2A and B***). Other cell types tested, e.g., HeLa, CHO, MDCK or cortical astrocytes displayed similar responses (***Figure 2B-figure supplement 4).*** We tested the effect of Bafilomycin A1. Bafilomycin A1 (1 µM) added acutely or following incubation for 1 h had little effect on the [Ca^2+^]_EL_ (***Figure supplement 5***), consistent with the failure to reconstitute the probe in its presence.

**Figure 2.**
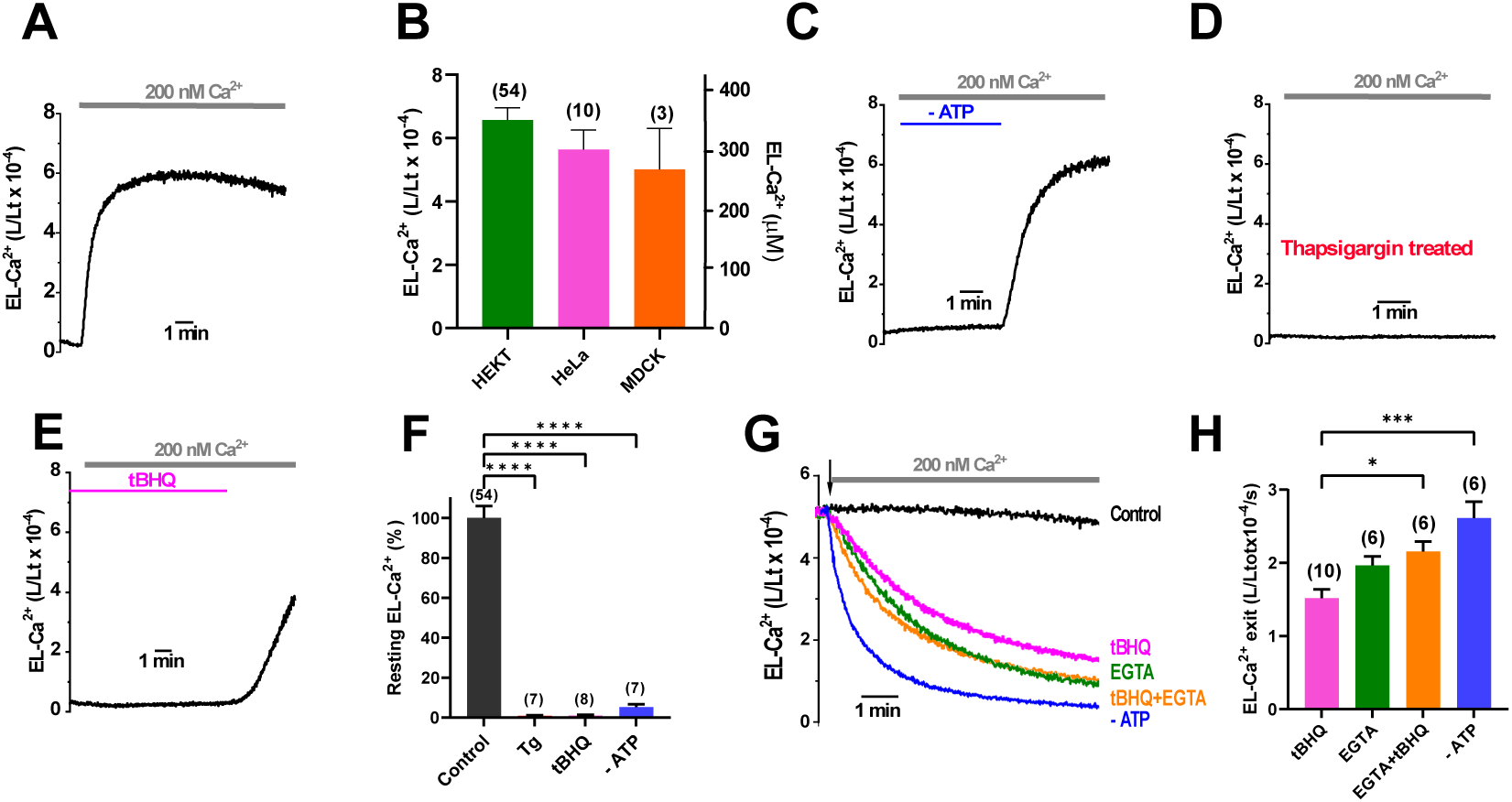
The endo-lysosomal Ca^2+^ store accumulates Ca^2+^ via SERCA. **(A)** Representative EL-Ca^2+^ signals recorded in HEK293T cells transiently transfected with ELGA1. After Ca^2+^ depletion prior to reconstitution with coelenterazine *n*, cells were permeabilized in Ca^2+^-free intracellular medium (containing 1 mM ATP) and then perfused with 200 nM free [Ca^2+^], as indicated. **(B)** Summary data (mean ± S.E.M.) of [Ca^2+^]_EL_ at steady-state obtained in populations of HEK293T, HeLa and MDCK cells with the number of independent transfections in parenthesis. **(C)** Cells were perfused as in (A) with an intracellular medium without ATP (-ATP) and reintroduced (1 mM) when indicated. **(D)** Cells were incubated with thapsigargin (1 µM) during the final 10 min of reconstitution followed by the application of the identical protocol described in (A). **(E)** Cells were perfused in an intracellular medium containing 2,5-di-tert-butylhydroquinone (tBHQ, 10 µM) followed by washout of the inhibitor. **(F)** Summary results (mean ± S.E.M.) with statistical analysis. ANOVA with Tukey’s post hoc test was applied. **(G)** Cells were perfused as in (A) and, once the [Ca^2+^]_EL_ steady-state was reached, different conditions were tested (arrow): none (200 nM Ca^2+^; Control), EGTA (0.1 mM), tBHQ (10 µM), EGTA+ tBHQ, or removal of ATP (-ATP)**. (H)** Summary results of EL-Ca^2+^ exit responses shown in (G), obtained from at least 3 independent days. ANOVA with Tukey’s post hoc test was applied. *p <D0.05, ***p <D0.001, and ****p <D0.0001.

Ca^2+^ uptake was ATP-dependent because there was little change in luminescence in the absence of ATP (90 ± 4% inhibition, n=7; ***Figure 2C and F-figure supplement 1B-C*).** Similarly, the SERCA specific inhibitors thapsigargin or tBHQ provoked a complete inhibition of the EL Ca^2+^ uptake (98 ± 1%, n=7; and 97 ± 2%, n=8; ***Figure 2D and E,*** respectively). In this last case, washing out of the reversible inhibitor rapidly restored the Ca^2+^ uptake up to normal rate (***Figure 2E***). Once the endo-lysosomal lumen was fully replenished with Ca^2+^, its removal (addition of 0.1 mM EGTA) caused a progressive and complete decline in the [Ca^2+^]_EL_ at a rate of ∼2.0 µM/s (***Figure 2G)***. Likewise, removal of ATP or application of tBHQ, either in the presence or absence of Ca^2+^, emptied the EL-Ca^2+^ store. Statistical quantifications of the EL-Ca^2+^-uptake and Ca^2+^-exit rate are shown in ***Figure 2F and H***.

Collectively, these results indicate that the endo-lysosome can accumulate high amounts of Ca^2+^ through a SERCA mechanism.

### IP_3_ mobilizes Ca^2+^ from endo-lysosomal Ca^2+^ stores

To explore endogenous Ca^2+^ release pathways, we examined the effect of NAADP, which activates endo-lysosomal TPCs. NAADP evoked a fast reversible response that released 30% of the EL-Ca^2+^ content, both in HEK and HeLa cells. But the responses were inconsistent and observable in around ∼ 50% of trials (***Figure supplement 6***). Surprisingly, addition of IP_3_ (2 µM, 30s) reliably produced a fast and strong Ca^2+^ release from the endo-lysosomal store that reduced the endo-lysosomal luminal [Ca^2+^]_EL_ by 72 ± 1% (n= 18) **(*Figure 3A and 3F*)**. The IP_3_-evoked Ca^2+^ release was very robust and consistent in all cell types tested (HeLa, CHO or MDCK cells; representative results are shown in ***figure supplement 7***). This effect was concentration dependent, from 0.01 µM, that produced a small but detectable effect, up to 10 µM, a maximal effect, comparable to that with 30 µM, the highest IP_3_ concentration tested **(*Figure 3B,*** and summary data in ***Figure 3C*)**.

**Figure 3.**
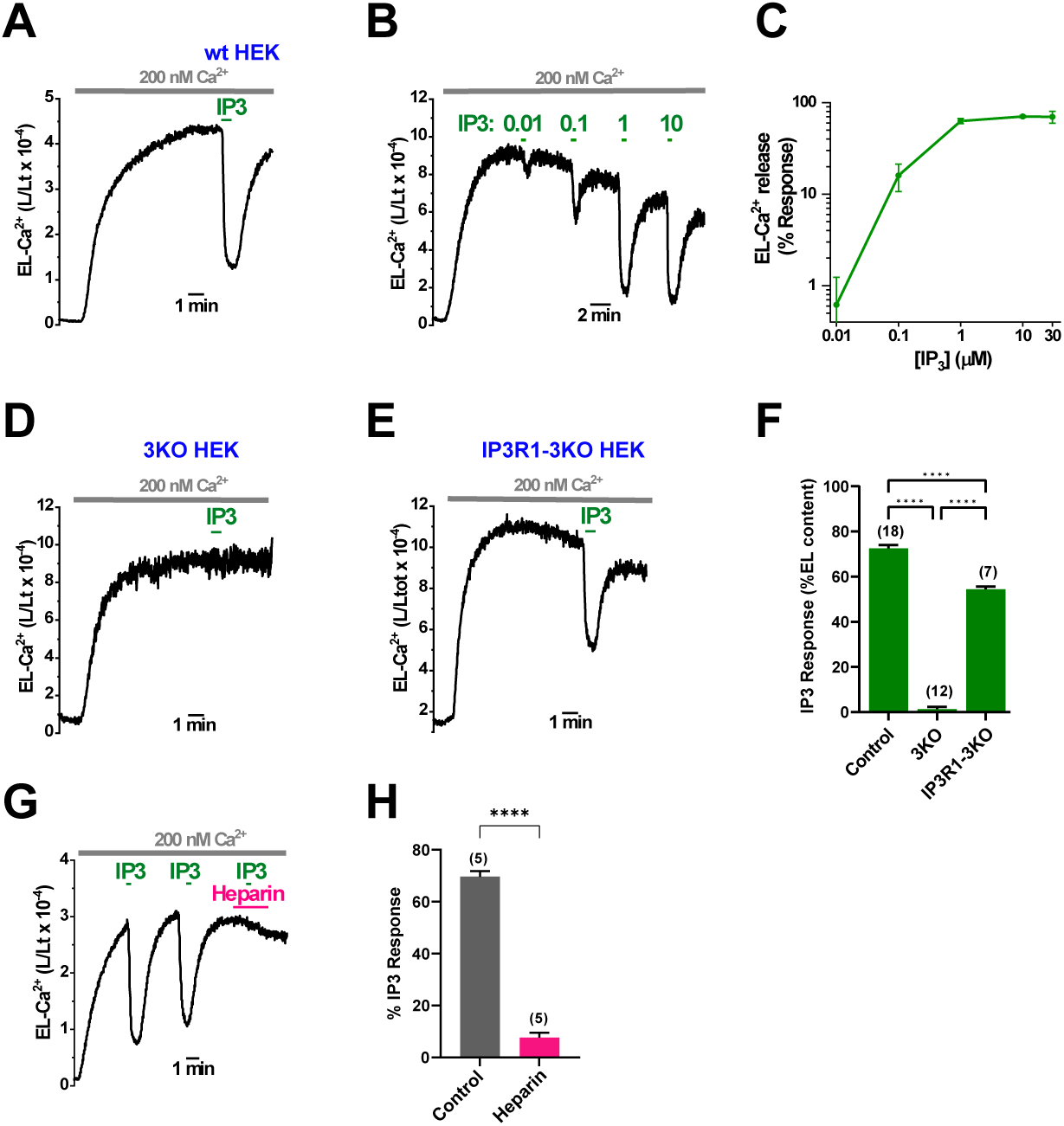
IP_3_ releases Ca^2+^ from the endo-lysosomal Ca^2+^ store. **(A)** Representative response to IP_3_ (2 µM, 30 s) in wt HEK293T cells expressing ELGA1. **(B)** Representative EL-Ca^2+^ release (in L/Lt ratio) upon application of four consecutive pulses (30s) of increasing concentrations of IP_3_ (in µM). **(C)** Concentration-dependent response of EL-Ca^2+^ release (expressed as % of maximal response obtained with 30 µM IP_3_) upon addition of increasing concentrations of IP_3_ (0.01, 0.1, 1 and 10 µM). Each point is the mean ± S.E.M. of 3-4 values obtained from independent transfections. **(D)** Representative response to IP_3_ (2 µM, 30 s) in IP_3_R-null HEK293 (IP3R-3KO) cell line expressing ELGA1. **(E)** Representative response to IP_3_ in IP3R-3KO cells transiently expressing IP_3_R1. **(F)** Statistical quantification of IP_3_ induced-EL-Ca^2+^ release (mean ± S.E.M.) with the number of independent transfections in parenthesis. ANOVA with Tukey’s post hoc test was applied. **(G)** Inhibition of the IP_3_ EL-Ca^2+^ release by heparin (100 µg/mL) in ELGA1 expressing HEK293T cells. **(H)** Statistical quantification of heparin inhibition of IP_3_ response. Data are expressed as mean ± S.E.M. In all bar graphs, numbers in parenthesis correspond to the number of independent transfections from at least 2-3 experiments. A Student′s *t* test was applied. ****p <D0.0001.

To test specificity of the IP_3_ response, we used IP_3_R-null HEK293 (HEK-3KO) cell line generated by CRISPR/Cas9 technology lacking all three IP_3_ receptor subtypes (Alzayady et al., 2016). As shown in ***Figure 3D***, IP_3_-induced Ca^2+^ release was abrogated (1.36 ± 0.97 % (n= 12). Interestingly, in these cells both the EL-Ca^2+^ refilling rate (6.7 ± 0.7 µM/s) and the [Ca^2+^]_EL_ reached at the steady-state (369 ± 32 µM; n= 12) coincided with those values obtained in IP_3_ receptor replete control cells (8.0 ± 0.6 µM/s; n= 36 and 365 ± 21 µM; n= 54) (***figure supplement 8)***. This suggests that, unless there are compensatory effects, IP_3_R is not essential for refilling the EL Ca^2+^ store. We also examined the effect of IP_3_ in HEK-3KO cells upon re-expression of IP_3_R1. As shown in ***Figure 3E and F***, the IP_3_ response was partially rescued (54 ± 1 %; n= 7).

In an independent approach, we used heparin, an inhibitor of the IP_3_R. Heparin completely blocked the IP_3_ response (92 ± 2 % inhibition; n= 5; ***Figure 3G and 3H*)**. Collectively, both genetic and pharmacological approaches reveal the presence of functional IP_3_Rs within the EL Ca^2+^ store that support robust luminal Ca^2+^ responses.

### ELGA responses are independent of the ER

Because of the similarities in the properties of the Ca^2+^ stores reported by ELGA and the ER with respect to both Ca^2+^ uptake and Ca^2+^ release, we considered the possibility that a fraction of the probe may be mistargeted to the ER.

To address this, we expressed TRPML1, a major endo-lysosomal Ca^2+^ channel. Colocalization analysis of ELGA with TRPML in fixed cells is shown in ***Figure 4A***. Both probes were targeted to the same compartment. Colocalization significantly decreased in live cells (***Figures 4A and B***) indicating that a proportion of ELGA is located to acidic, non-fluorescent vesicles in live cells that become fluorescent when the pH increases upon fixation. These results provide further evidence that ELGA is a Ca^2+^ reporter in a fraction of the EL system.

**Figure 4.**
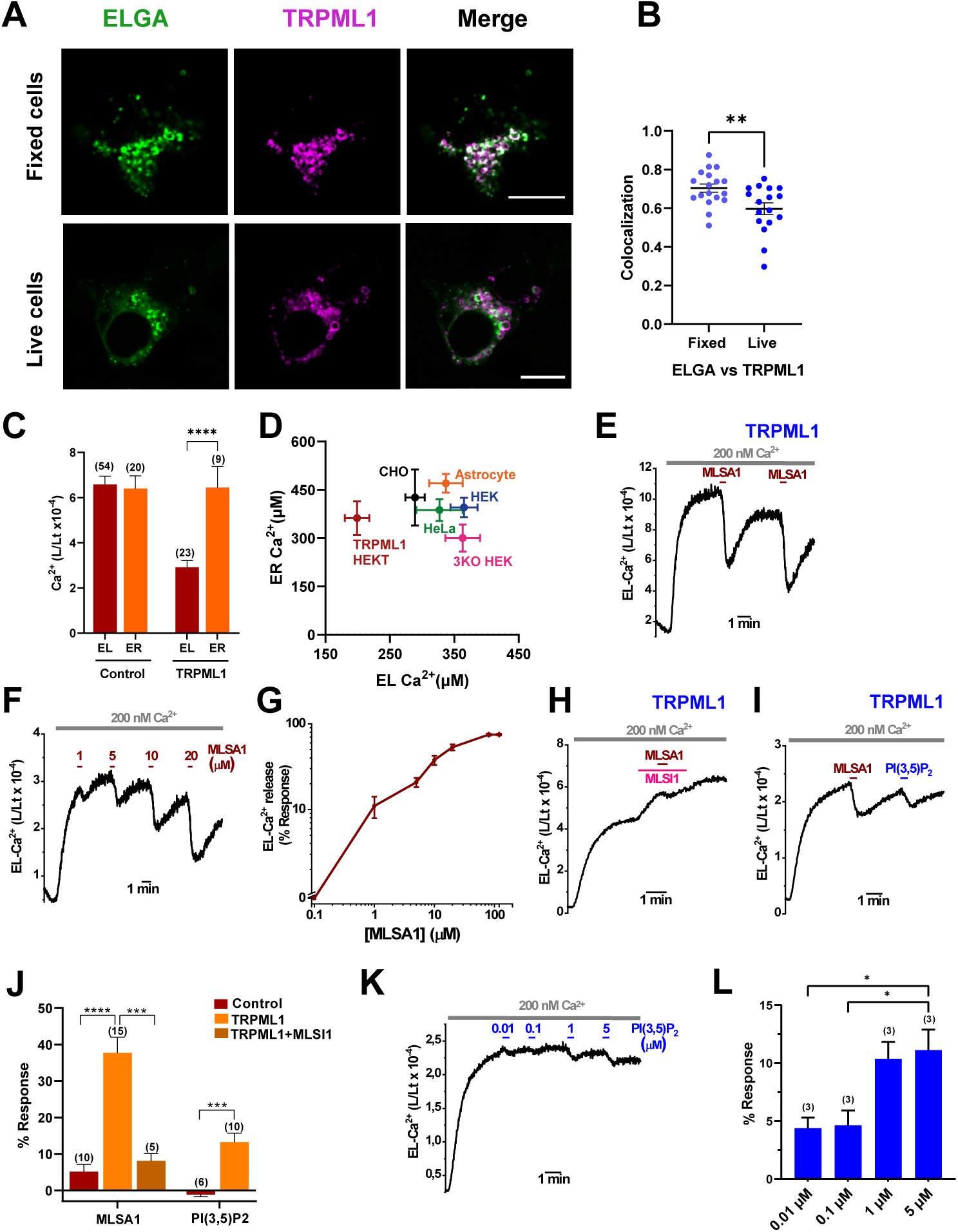
ELGA is functionally located to the endo-lysosome. **(A)** Colocalization of ELGA fluorescence with TRPML1-cherry (magenta) in either fixed or live HEK293T cells expressing both proteins. Scale bar in all images, 10 µm. **(B)** Quantitative colocalization analysis of confocal fluorescence microscopy images were expressed as Pearson’s (*P*) correlation coefficient shown in (A). Each point represents an individual cell from at least 2 independent experiments (mean ± S.E.M). A Student’s *t* test was applied. **(C)** Effect of expression of TRPML1 on the [Ca^2+^] at steady-state recorded from HEK293T expressing either ELGA1 (EL) or ERGA1 (ER). Data are expressed as mean ± S.E.M. In all bar graphs, numbers in parenthesis correspond to the independent transfections. A Student’s *t* test was applied. **(D)** Lack of correlation between resting [Ca^2+^] in the ER and the EL recorded with ERGA1 or ELGA1, respectively, in six cell types: HEK, TRPML1-expressing HEK, 3KO HEK, HeLa, CHO cells and cultured murine cortical astrocytes. Data are expressed as mean ± S.E.M. ER-Ca^2+^ in astrocytes taken from (Rodríguez-Prados et al., 2020). **(E)** Representative ML-SA1 (20 µM, 30 s) responses inducing EL-Ca^2+^ releases (expressed as L/Lt) from HEK293T cells expressing TRPML1 measured with ELGA1. **(F)** Representative Ca^2+^ responses (expressed as L/Lt) to application of four consecutive pulses (30s) of increasing concentrations of ML-SA1. **(G)** Concentration-dependent EL-Ca^2+^ release (expressed as % of the maximal response) in response to increasing concentrations of ML-SA1. Each point is the mean ± S.E.M. of 3-4 values obtained from 3 independent experiments. **(H)** Effect of the TRPML1 inhibitor ML-SI1 (10 µM) on the ML-SA1 (20 µM) response (expressed as L/Lt) in TRPML1 transfected HEK293T cells. **(I)** Effect of PI(3,5)P_2_ (1 µM) on the EL-Ca^2+^ release in TRPML1 expressing HEK293T cells. **(J)** Summary results and statistical quantification of EL-Ca^2+^ release responses under various experimental conditions. Control (non-transfected) cells and TRPML1 transfected HEK293T cells were stimulated with ML-SA1 or PI(3,5)P_2_. Percentage of response related to total [Ca^2+^]_EL_ at the steady-state is indicated. Data are expressed as mean ± S.E.M. and numbers in parenthesis indicate total *n* of independent transfections. One-way ANOVA with a Tukey’s post hoc test (MLSA1 data) or a Student’s *t* test (PI(3,5)P_2_) were applied. **(K)** Concentration-dependent response (expressed as L/Lt) for EL-Ca^2+^ release upon increasing concentrations of PI(3,5)P_2_. **(L)** Summary results and statistical significance of the responses determined from three independent transfection are shown in (K). Results are expressed as mean ± S.E.M. An ANOVA with a Tukey’s post hoc test was applied. ****p <D0.0001 and ***p <D0.001.

Interestingly, TRPML1 expression drastically reduced the levels of Ca^2+^ reported by ELGA1 (***Figure 4C)***. In contrast, Ca^2+^ levels in the ER measured using the same reporter targeted to the ER were unchanged upon TRPML1 expression, indicating that the channel is specifically leaky in the EL store. These functional data distinguish EL and ER Ca^2+^ stores. In accord, there was no correlation between EL and ER Ca^2+^ levels reported by the corresponding GA1 in the cell types used (***Figure 4D)*.**

To further explore the properties of TRPML1, we activated it with the specific membrane-permeable synthetic agonist ML-SA1. In transiently expressing TRPML1 HEK293T cells, ML-SA1 (20 µM, 30s) produced a robust EL-Ca^2+^ release of 38 ± 4% (n= 15) of the total EL luminal Ca^2+^content, as reported by ELGA1 at the steady-state (***Figure 4E and J***). Upon application, ML-SA1 exhibited a lag time of ∼20 s and it continued releasing Ca^2+^ once the agonist was washed out. Consecutive applications of ML-SA1 evoked responses of similar amplitude, indicating no desensitization of the channel. The time to refill the store after the washing out was ∼5 min. Full concentration-effect relationship for the Ca^2+^ release response showed an EC_50_ value of 7 ± 1 µM (***Figures 4F and G***). Comparable responses were observed in TRPML1-expressing HeLa cells (***figure supplement 9)***. In accord, the response to ML-SA1 was selectively blocked by the specific TRPML-antagonists ML-SI1 **(*Figure 4H-figure supplement 9B and 9C*)** or ML-SI3 (***figure supplement 9C***) and expression of TRPML1 did not affect the response to IP_3_ (***figure supplement 9D*).**

We also activated TRPML1 with its endogenous activator PI(3,5)P_2_. As shown in ***Figure 4I***, PI(3,5)P_2_ evoked EL-Ca^2+^ release in cells expressing TRPML1 albeit modestly compared to ML-SA1. Statistical quantification is summarized in ***Figure 4J***. The EL-Ca^2+^ release provoked by PI(3,5)P_2_ was concentration-dependent (***Figure 4K and L)*.**

These data show that ELGA accurately reports the properties of a *bona fide* EL Ca^2+^ channel.

### A subset of IP_3_ receptors localize to endo-lysosomes

We applied several high resolution microscopy techniques to assess whether IP_3_ receptors localized to the EL system. We used HeLa cells in which endogenous IP_3_R1 is tagged with EGFP by gene editing (Thillaiappan et al., 2017). Results from stimulated emission depletion (STED) super-resolution microscopy showed that IP_3_R1-GFP displayed a well resolved punctate distribution in fixed cells (***Figure 5A***). This distribution was compared to expressed EL channels. We detected a number of IP_3_R1 puncta that colocalized with TRPML1 (***Figures 5Aa-c***). Essentially, similar results were obtained with TPC2 (***Figures 5Ad-f***). In addition, we found part colocalization of IP_3_R1 (***Figures 5Ag-i***) and SERCA2 (***figure supplement 10***) with endogenous LAMP2 although overlap was modest. We also used STED microscopy to localize endogenous IP_3_R3 in HEK cells. We found evidence for part colocalization of IP_3_R3 with endogenous LAMP1 (***Figure 5B)***.

**Figure 5.**
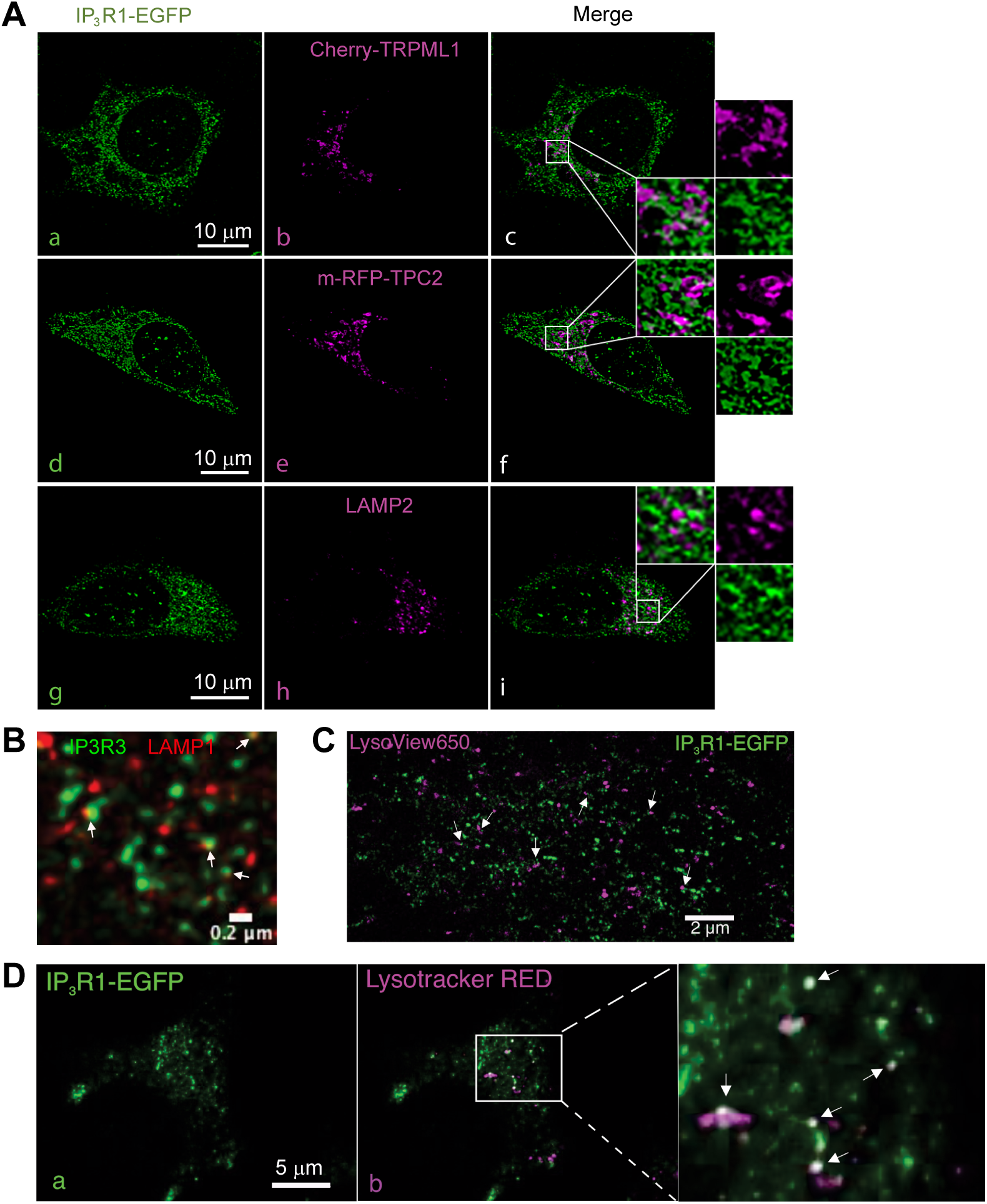
ELGA is physically located to the endo-lysosome. **(A)** Colocalization of endogenous IP_3_R1 with various endo-lysosome markers in EGFP-IP_3_R1 HeLa cells using STED microscopy. Cells were transiently transfected with cherry-TRPML1 (a-c) or mRFP-TPC2 (d-f). EGFP-IP_3_R1 and LAMP2 were detected using specific antibodies against EGFP or LAMP2. **(B)** Association (arrows) of endogenous IP_3_R3 with LAMP1 detected with specific antibodies anti-IP_3_R3 and anti-LAMP1 in fixed wild type HEK293 cells using STED Super-resolution microscopy. **(C)** Colocalization (arrows) of endo-lysosomes stained with LysoView650 (pink) and endogenous IP_3_R1-EGFP in live Hela cells using live cell STED microscopy. **(D)** Colocalization (arrows) of endo-lysosomes stained with lysotracker red (pink) and endogenous IP_3_R1-EGFP in HeLa cells using TIRFM.

To seek evidence for IP_3_R localization with EL system in live cells, we loaded IP_3_R1-GFP HeLa cells with fluorescent acidotropes (LysoView650 and Lysotracker red). Some IP_3_R1 clearly formed puncta with endo-lysosomes using either super-resolution microscopy (STED; ***Figure 5C)*** or total internal reflection fluorescence microscopy (TIRFM; ***Figure 5D)*.**

In sum, these data support the view that a subpopulation of IP_3_Rs are physically localized to the endo-lysosomal system.

### Physiological IP_3_-forming stimuli mobilize Ca^2+^ from endo-lysosomal Ca^2+^ stores

To further characterize endo-lysosomal Ca^2+^ dynamics, we assessed luminal Ca^2+^ levels in intact cells. Addition of 1 mM CaCl_2_ refilled the EL Ca^2+^store up to a steady-state of 234 ± 29 μM (n= 12) (***Figure 6A***).

**Figure 6.**
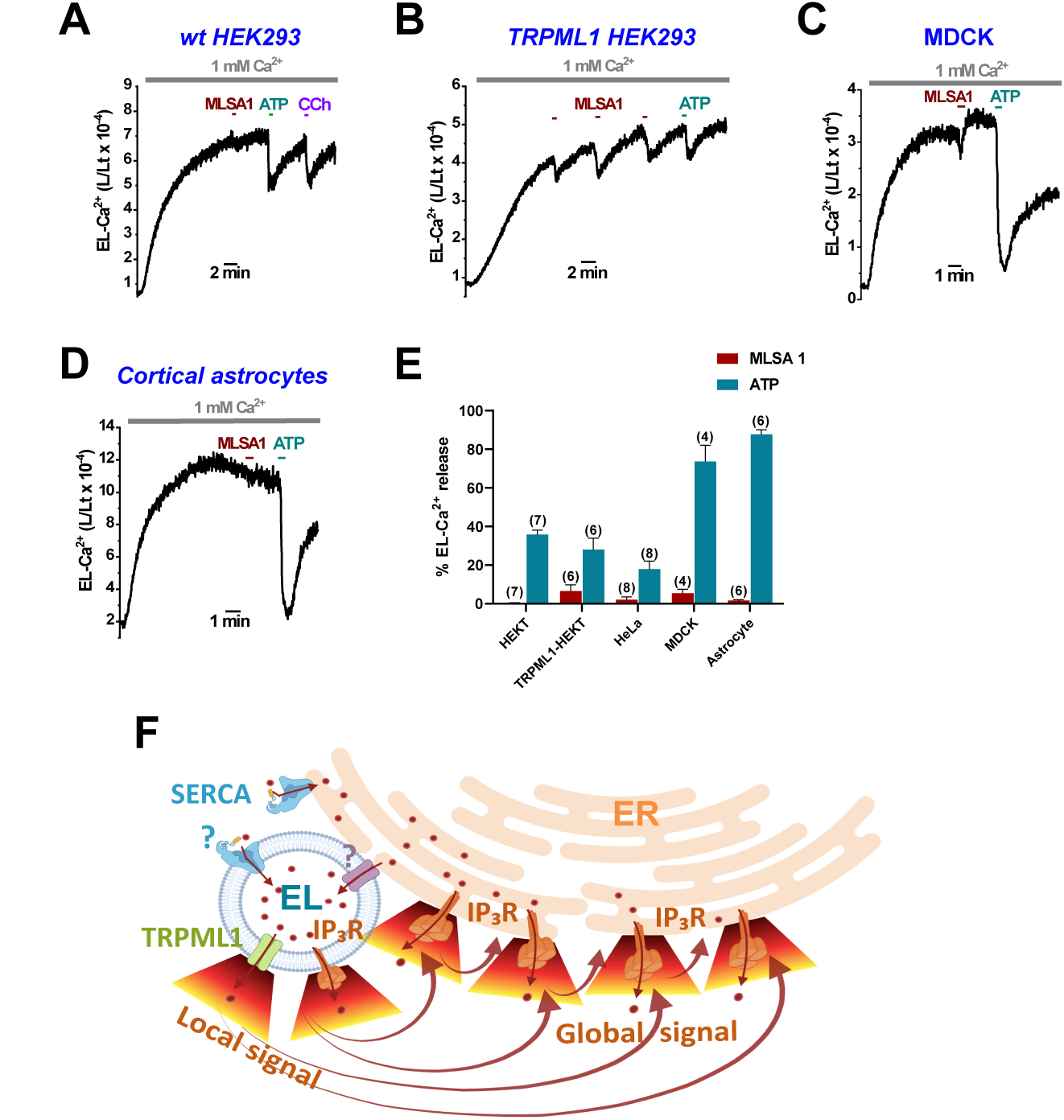
Physiological endo-lysosomal Ca^2+^ release. **(A)** Representative EL-Ca^2+^ responses (in L/Lt) to agonists (ML-SA1, 20 µM; ATP, 100 µM; Carbachol, (CCh), 100 µM) in intact HEK293T cells expressing ELGA1. **(B)** Similar to (A), in transiently expressing TRPML1 HEK293T cells. **(C)** Protocol similar to (A) applied in MDCK cells. **(D)** Protocol similar to (A) applied in astrocytes. **(E)** Summary results of EL-Ca^2+^ release induced by ML-SA1 and ATP in different cell types or conditions. Data are expressed as mean ± S.E.M. and numbers in brackets correspond to the total number of independent transfections. Responses are in percentages related to total [Ca^2+^]_EL_ at the steady-state. **(F)** A proposed model of Ca^2+^ dynamics in the endo-lysosome (see text for explanation).

To probe the functionality of IP_3_ receptors, we stimulated live cells with ATP which activates endogenous P2Y receptors or carbachol which activates muscarinic receptors. ATP and carbachol provoked fast and reversible EL Ca^2+^responses of 35.81 ± 2.34 % (n= 7) and 19.88 ± 1.87 % (n= 5), respectively (***Figure 6A***). ML-SA1 also produced reproducible responses in cells expressing TRPML1 although it was ineffective in wild type cells (***Figure 6A and B***). Similar experiments were performed in polarised MDCK cells ***(Figure 6C).*** This cell type displayed a comparable steady-state EL-Ca^2+^ that was almost completely discharged upon cell activation with ATP (73.72 ± 16.85 %; n= 4).

Finally, we examined EL responses in a primary cell type. In mouse cortical astrocytes ***(Figure 6D)***, ML-SA1 application elicited a negligible effect whereas a subsequent pulse of ATP almost completely discharged the EL-Ca^2+^ store (87.68 ± 5.90 %; n= 6). A summary of the data is shown in ***Figure 6E***.

In sum, these data identify a widespread physiologically-relevant IP_3_-sensitive Ca^2+^ store within the EL system.

## DISCUSSION

Using a new genetically-encoded Ca^2+^ indicator based on aequorin, we demonstrate here the existence of a *bona-fide* endo-lysosomal Ca^2+^ store, able to accumulate a high [Ca^2+^] and to release it upon physiological activation (***Figure 6F***).

Our results are best explained if this endo-lysosome is functionally coupled with the ER to accumulate Ca^2+^ via active transport such as via SERCA pumps. Although the two compartments work in parallel, they are distinct in the amount of Ca^2+^ that they can store, with the luminal Ca^2+^ concentration being significantly lower in the EL system (∼200-400 µM) than the ER (∼400-500 µM) and in their sensitivity to expression of TRPML1, which selectively lowers the EL Ca^2+^ store content. The EL Ca^2+^ values are in agreement with previous reports (Narayanaswamy et al., 2019; Ronco et al., 2015) but contrast with others (Albrecht et al., 2015). The SERCA might be located at the EL membrane where it directly pumps Ca^2+^ into its lumen, or, alternatively, Ca^2+^ could be indirectly fuelled from the ER into the EL lumen, through an undetermined channel connecting the two organelles. The ER directly contacts the EL system through membrane contact sites (MCS), especially abundant in late endosomes, where they allow exchange of lipids and other molecules, including Ca^2+^ (Kilpatrick et al., 2013; Lee and Blackstone, 2020; Raiborg and Stenmark, 2015). Close association of ELGA-positive compartments with ER reported here might provide a route for uptake.

We show that activation of TRPML1 with ML-SA1/PI(3,5)P_2_ and TPCs with NAADP produces luminal responses, but the latter were not consistent. Recent studies have shown that NAADP binds to soluble binding proteins to activate TPCs (Gunaratne et al., 2023; Saito et al., 2023). It is possible that loss of these proteins upon permeabilization may account for NAADP insensitivity. Surprisingly, IP_3_ consistently produced robust Ca^2+^ release responses comparable to those evoked by bonafide EL channels. In accord with these functional data, our results with super resolution microscopy and TIRFM provide direct evidence for the presence of IP_3_R on the endo-lysosome vesicles. Importantly, proteomic analysis of endo-lysosomal vesicles isolated from HEK293 cells by multiple different protocols, has also identified the two isoforms of IP_3_R and SERCA in this organelle (Singh et al., 2020). Moreover, IP_3_Rs are detected in numerous additional cell lines, both of mouse and human origin (Akter et al., 2023). Finally, we have also confirmed the presence of IP_3_Rs and SERCA in endo-lysosomes using LysoIP (unpublished results). Functional IP_3_ receptors have been documented in other acidic organelles such as secretory granules in professional secretory cell types (Gerasimenko et al., 2006; Santodomingo et al., 2008) and the acidocalcisome in Trypanosomes (Huang et al., 2013). But we are not aware of their presence within the EL system, which extends IP_3_ action to potentially all cells.

Previous work suggested that IP_3_ fills the EL system with Ca^2+^ (Atakpa et al., 2018; Garrity et al., 2016) as opposed to emptying it as reported here. But direct measurements of luminal Ca^2+^ in response to IP_3_ were not reported previously. Instead, these conclusions were based almost exclusively on indirect measurements of Ca^2+^ in the cytosol. Additionally, because reconstitution of ELGA is pH sensitive, it reports Ca^2+^ in only a sub-compartment of the EL system. The EL system is a hybrid compartment, very heterogeneous in morphology, composition, and function. Although this organelle is globally considered acidic, its luminal pH varies from 4.5 up to 6.5 in different vesicles. The acidity of the vesicles has been correlated with subcellular location (Bright et al., 2016; Johnson et al., 2016) likely indicative of functional diversity. Thus, Ca^2+^ handling, might also show heterogeneity among the various endo-lysosomal vesicles not least because of the relationship between luminal Ca^2+^ and pH (Christensen et al., 2002). Probes that allow spatial mapping of Ca^2+^ are required to pinpoint the presence of endosomal IP_3_ receptors that might release Ca^2+^ and ER IP_3_ receptors that might drive Ca^2+^ uptake.

Our intact cell studies provide evidence that IP_3_-sensitive stores can be tapped during physiological stimulation with IP_3_-forming stimuli. The fast exit of Ca^2+^ from the EL into the cytosol through IP_3_Rs would generate local cytosolic Ca^2+^ signals. These, in turn, could be amplified either via Ca^2+^-induced Ca^2+^ release (CICR) through IP_3_Rs (or RyRs in other cell types) present in the ER membrane to generate global signals. Indeed, IP_3_Rs have been shown to be preferentially associate with the lysosomes at contact sites (Atakpa et al., 2019). Such coupling would be analogous to that triggering of Ca^2+^ signals by TRPML (Kilpatrick et al., 2016) or TPCs (Xu and Ren, 2015; Yuan et al., 2024; Zhu et al., 2010). Our results are in line with the ‘*chatter’* between the two organelles (Patel et al., 2010) and the essential role of the ER in the EL signal in various cell types (Garrity et al., 2016; López-Sanjurjo et al., 2013; Morgan et al., 2013; Yang et al., 2019). Alternatively, local IP_3_R-dependent Ca^2+^ signals may regulate endo-lysosomal function at basal (unstimulated) levels of IP_3_, again in analogy with TPCs and TRPML, which are well established in regulating membrane traffic (Pryor et al., 2000; Ruas et al., 2010; Vassileva et al., 2020).

In conclusion, our direct luminal Ca^2+^ measurements help to fill the gaps in our knowledge on the Ca^2+^ signalling toolkit of the endo-lysosome system and provide new insight into the dynamics of these small acidic Ca^2+^ stores and their functional relationship with the ER Ca^2+^ stores.

## METHODS

### Gene construction

*H. sapiens* VAMP7 cDNA was amplified by PCR using specific primers (#324; forward: 5’-CCAaagcttGCCACCATGGCGATTCTTTTT-3’; #325; reverse: 5’-ATaagcttTTTCTTCACACAGCTTGGCCATG-3’; *HindIII* sites are in lowercase). This fragment was digested with *HindIII* and inserted upstream of the *HindIII* fragment encoding the GA or the GA1 cDNAs, in the pHSV plasmid leading to pHSV-ELGA or pHSV-ELGA1 plasmids, respectively. The GA gene is a fusion of EGFP and aequorin (Manjarres et al., 2008) containing the D119A mutation (Kendall et al., 1992). The GA1 (previously described as GAP1) gene is a fusion of GFP (C3) and aequorin containing the mutations D117A, D119A and D163A (Rodriguez-Garcia et al., 2014). For expression in bacteria, GA or GA1 cDNAs were cloned into the pET28a vector. For measurements of ER-Ca^2+^, the pHSV-ERGA1 (previously described as erGAP1) was used (Rodriguez-Prados et al., 2015).

### Virus production

pHSV-ELGA or pHSV-ELGA1 were packaged into HSV-1 amplicon particles using a deletion mutant packaging system. Briefly, these plasmids were transfected into 2-2 cells and then, infected with a defective 5dl 1.2 helper virus. Titration was performed as previously described (Alonso et al., 1996; Lim et al., 1996).

### Bacterial expression of GA and GA1 proteins

GA or GA1 cloned in pET28a were transformed in *E. coli* BL21 (Stratagene). Bacteria were grown at 37° C to A_600_ >0.6 in LB containing 40 mg/L kanamycin and induced with 0.05 mM isopropyl β-D-thiogalactoside (IPTG) for 5h at 30°. Cells were then pelleted by centrifugation at 6000 g for 30 min and sonicated in a buffer containing (in mM): NaCl, 50; DTT, 0.5; Tris-HCl 20, pH 7.5; and Roche protein inhibitors. The bacterial lysate was centrifuged at 30,000 g for 10 min. Protein was purified by using a HisPur Ni-NTA resin (ThermoFisher, 88222). Protein quantification was performed by Bradford (range 20-60 µg/µL).

### Bioluminescence activity at various pHs

Measurements were performed in the Tecan Genios Pro 96-well plate reader. Recombinant *E.coli* GA or GA1 protein was reconstituted with coelenterazine *n* (1 µM) for 20 min and then 5-10 µL were aliquoted into each well to a final volume of 100 µL with a buffer containing (in mM): KCl, 140; MgCl_2_, 1; MOPS/Tris, 20, pH 7.2. The effects of pH were monitored using different buffers for each pH range: acetate for pH in the range 4.5-5.5; MOPS/Na for pH range 6-6.5; and Hepes/Na for pH 7.2. Maximal luminescence was acquired by injecting 10-25 µL of saturating (50 mM) Ca^2+^. Calibrations were performed in Ca^2+^-free medium (0.1 mM EGTA added), or in the same solutions containing increasing Ca^2+^ concentrations between 50 µM and 50 mM. This Ca^2+^solution (10 µL) was injected with a first injector, read for 15 seconds and then a saturating Ca^2+^ solution (10 mM) was injected with a second injector.

### Cell culture and gene expression

HeLa (CCL-2), HEK 293-T (CLR-1573), CHO-K1 (CCL-61) or MDCK cells (CCL-34), were all purchased from ATCC. EGFP-IP_3_R1 HeLa cells were earlier described (Thillaiappan et al., 2017). All cells were maintained in Dulbecco’s Modified Eagle Medium (DMEM, Invitrogen) supplemented with 10% fetal bovine serum (FBS, Sigma), 2 mM L-glutamine, 100 µg/ml streptomycin and 100 U/ml penicillin. IP_3_R-null HEK293 (HEK-3KO) cells (Alzayady et al., 2016) were maintained in DMEM/F-12 with GlutaMAX (ThermoFisher) supplemented with 10%, FBS. Periodic PCR screening confirmed that all cells were free of mycoplasma infection.

Mouse cortical astrocytes were isolated using the protocol described previously (Rodríguez-Prados et al., 2020) with the approval of the Institutional Animal Care and Use Committee at the University of Valladolid (ethics approval reference number: 10207729). Briefly, cortices were dissected from three P2-3 C57BL/6J Crl (Charles Rivers) mice in cold HBSS and digested with 0.5 mg/mL papain and 0.04 mg/mL DNase dissolved in a Ca^2+^-and Mg-free HBSS with 10 mM glucose and 0.1% bovine serum albumin (BSA) at 37°C for 20 min. Cells were resuspended with Minimal Essential Medium (MEM) supplemented with 2 mM glutamax, 27 mM glucose, 100 U/mL penicillin, 100 µg/mL streptomycin and 10% (v/v) horse serum, and plated on a poly-L-lysine coated 75 cm^2^ flask. After 2 and 7 days, flasks were slapped to remove neurons and other loosely attached cells such as microglia and oligodendrocytes. After one week, cells were trypsinized and seeded in the same medium onto poly-L-lysine coated 12 mm-diameter glass coverslips at a density of 2.5 x 10^4^ cells/coverslip. The cultures were used within 1-2 weeks.

Cells were transiently transfected with 0.2 µg (HEK293T) or 0.2-0.5 µg (HeLa) pHSV-ELGA or pHSV-ELGA1 using Lipofectamine 2000 (Invitrogen), according to the manufacturer’s instructions. In some experiments, this plasmid was cotransfected with other plasmids encoding: mRFP-TPC2 (0.1 µg); EGFP-trpML1 (0.02 µg); mCherry-trpML1 (0.05 µg; Addgene #135189); er-Ruby-GCaMP210 (0.5 µg); or IP_3_R1 (0.02 µg). Stably transfected HeLa clones expressing ELGA were generated by cotransfection of pHSV-ELGA and pcDNA3 (carrying the *neomycin* gene). GFP positive cells were enriched by sorting in a FACSAria II followed by antibiotic selection with 800 µg/mL G418 and single cell cloning was performed by limited dilution. ELGA-HeLa clones (No. 3 and 6) were maintained with 200 µg/mL G418.

ELGAs expression in mouse cortical astrocyte cultures was achieved by infection with herpes HSV-ELGA, HSV-ELGA1 or HSV-ERGA1 (previously named erGAP1) amplicons at a multiplicity of infection ranging between 0.01 and 0.1, one day before use.

### Ca^2+^ bioluminescence measurements

Cells expressing ELGA1 (or ELGA in some experiments) were seeded on 4-well plates at a density of 8 × 10^4^ (HeLa, CHO, or MDCK cells) or 2 × 10^5^ (HEK-293T and HEK-3KO cells). Cells were Ca^2+^ depleted by incubation with the reversible sarco-endoplasmic reticulum Ca^2+^-ATPase (SERCA) inhibitor 2,5-di-tert-butylhydroquinone (TBH; 10 µM) in the extracellular buffer (145 mM NaCl; 5 mM KCl; 1 mM MgCl_2_; 10 mM glucose; 10 mM Na-HEPES, pH 7.4) supplemented with 0.5 mM EGTA for 10 min at 22°C. Apo-aequorin was then reconstituted by incubation with 1 µM *n* coelenterazine in the same medium during 1 hour prior to measurements. Cells were then placed in the luminometer (Cairn Research, UK). Light acquisition was recorded each second as cps.

Intact cells were perfused (5 mL/min) with the extracellular buffer containing 1 mM CaCl_2_ plus additions as stated. In the experiments with permeabilized cells, perfusion was performed with an intracellular-like (IL) medium with the following composition (in mM): KCl, 140; KH_2_PO_4_, 1; MgCl_2_ 1; Mg-ATP, 1; sodium succinate, 2; sodium-pyruvate, 1; Na-HEPES, 20, pH 7.2. The cells were permeabilized by perfusion of digitonin (50 µM) in the IL medium containing 0.5 mM EGTA during 1 min. The digitonin was washed out and the cells were then switched to IL medium containing 200 nM Ca^2+^made by blending titrated solutions of EGTA and EGTA-Ca^2+^ in the required amounts, calculated using the program MaxChelator (Patton et al., 2004). If not stated otherwise, all stimuli or drugs were dissolved and applied in the IL-buffer containing 200 nM Ca^2+^ during 30s, as indicated in the graphs. At the end of each run, cells were lysed with 100 µM digitonin in a 10 mM CaCl_2_ solution to release all the remaining aequorin luminescence. The EL-Ca^2+^ was expressed as luminescence at a certain time point (L) divided by total luminescence till that moment (L/Lt). All measurements were performed at 22°C. Calibration curve allowed to transform L/Lt into [Ca^2+^] (µM) (Rodriguez-Prados et al., 2015).

### Immunofluorescence

2-5 x 10^4^ HEK cells were seeded onto 12 mm-glass coverslips and fixed, either with cold 100% methanol or with 4% PFA in PBS for 20 min, followed by incubation with 10% goat serum, 0.2% saponin (or 1% goat serum with 0.1% Tween-20 in some experiments) in PBS for 20 min at 22 °C. The incubation with the primary antibody was performed overnight at 4°C. The following primary antibodies were used: anti-IP3R3 (1:100; BD biosciences); anti-LAMP1 (1:500; Cell Signaling); anti-LAMP2 (H4B4) antibody (1:100; DSHB / University of Iowa); anti-EEA1 (1:200; BD Transduction Laboratories); anti-TOM20 (1:100; Santa Cruz); anti-GFP (1:500; Invitrogen). The secondary antibodies (anti-mouse Alexa Fluor 568-conjugated antibodies) were incubated for 1h at 22 °C. Nuclei were stained with 4’,6-diamidino-2-phenylindole (DAPI). Cells were mounted with VectashieldR (Vector).

### Confocal microscopy

Intact or fixed cells were imaged under a 63X Plan-Apochromat (NA 1.4) oil immersion objective using a confocal microscope (SP5 Leica) equipped with a white laser. Images were analyzed with ImageJ software using the JACoP plugin (Bolte and Cordelières, 2006).

### STED microscopy

3D STED microscopy was performed using a Leica SP8-STED microscope equipped with an HC PL APO CS2 x100/1.4NA oil immersion objective. Fluorescence depletion was made by a white laser and using a LAS-X deconvolution software. Some experiments were performed with an Abberior Instruments Expert Line STED microscope equipped with an Olympus UPLSAPO ×100/1.4NA oil immersion objective. Sequential confocal and STED images were obtained following excitation of Alexa Fluor 594 and STAR RED by 594 and 640 nm lasers, respectively. Both fluorophores were depleted in three dimensions with a 775 nm pulsed STED laser. Z-stacks were obtained by collecting images at 50 nm intervals using the 3D STED mode. Rescue STED was employed to minimize the light dosage.

### TIRFM

Images were acquired using an Olympus IX81 inverted Total Internal Reflection Fluorescence Microscopy (TIRFM) equipped with oil-immersion PLAPO OTIRFM 60x objective lens/1.45 numerical aperture. Olympus CellSens Dimensions 2.3 (Build 189987) software was used for imaging. The cells were illuminated using 488 or 568 nm laser and the emitted fluorescence was collected through a band-pass filter by a Hamamatsu ORCA-Fusion CMOS camera. The angle of the excitation beam was adjusted to achieve TIRF with a penetration depth of ∼140 nm.

### Statistical analysis

Results are expressed as mean ± S.E.M., as indicated. Data were analyzed with the GraphPad Prism 9 software (San Diego, CA, USA). Statistical significance between two groups was analyzed by the unpaired Student’s *t* test, whereas more than two groups were compared using one-way analysis of variance (ANOVA) and Tukey post-test analysis. Differences were considered statistically significant when probability (*p*) values were less or equal than 0.05 (*), 0.01 (**), 0.001 (***) or 0.0001(****). Sample sizes (n) refer to independent cell wells from, at least, 3 different days. Individual traces show representative experiments.

## Credit authorship contribution statement

FJA made the original observations; BC, PT-V, AD-L, CR, FJA, JR-R, BMM and DIY performed investigation, formal analysis, and interpretation of data; DIY, VM, JG-S, SP and MTA provided conceptualization and funding acquisition; MTA and SP wrote original draft; all authors reviewed and edited the manuscript.

## Acknowledgments

We thank Dr. H. Xu for the VAMP7 plasmid; Dr. De Juan-Sanz for erRuby-GCaMP210; and, C. Taylor for IP_3_R1 cDNA and EGFP-IP_3_R1 HeLa cells. We also thank Dr. T. Schimmang for his valuable comments and suggestions. We also thank I. López and J. Fernández, as well as the Servicio de Microscopía, (C.I.C., Salamanca), for technical assistance.

