## Supplemental material for "Direct measurements of luminal Ca^2+^ with endo-lysosomal GFP-aequorin reveal functional IP_3_ receptors"

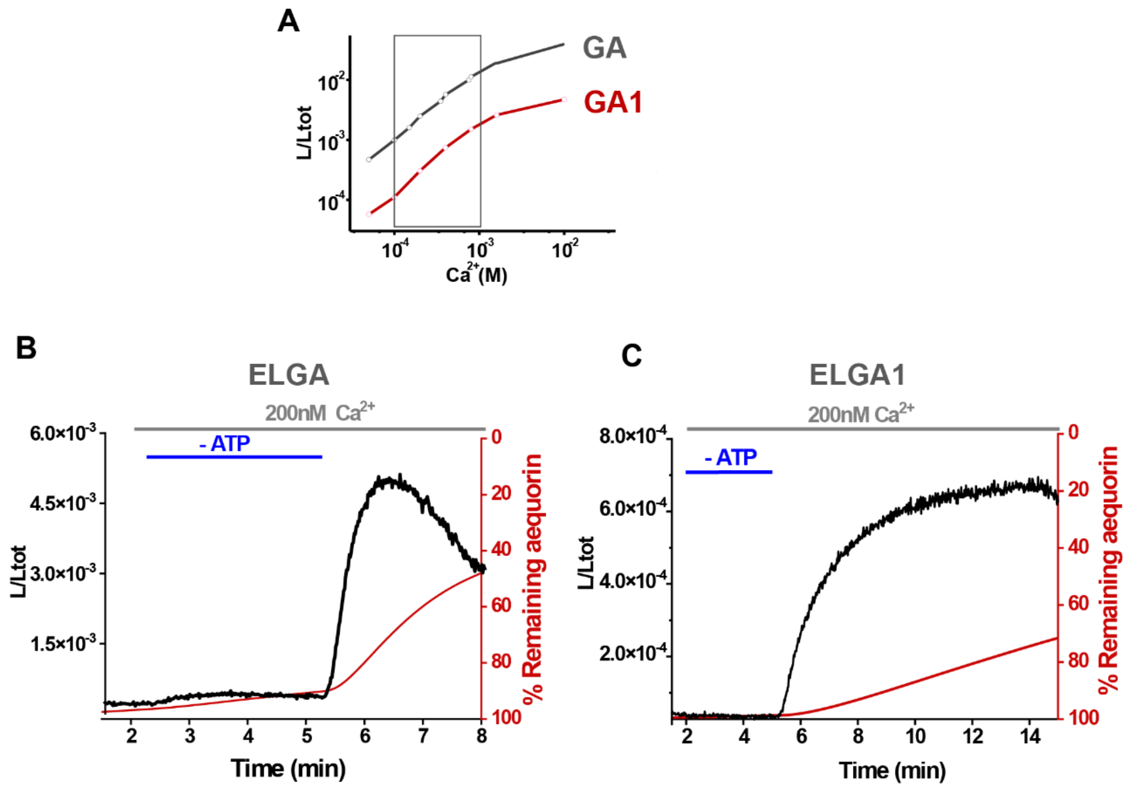


**Figure Supplement 1.**

**GA and GA1 bioluminescent $Ca^{2+}$ sensors. (A)** $Ca^{2+}$ calibration of GA and GA1. Marked region is the expected range of [$Ca^{2+}$] in the EL. **(B, C)** Comparison of luminescence emission and consumption (in % of the total) in a representative experiment in transiently transfected HeLa cells with ELGA **(B)** or ELGA1 **(C).** Notice the stability of the ELGA1 signal at the steady-state (less than 5% at min 8) in comparison with the ELGA signal that quickly declines due to the rapid consumption (~50% at the same timepoint).

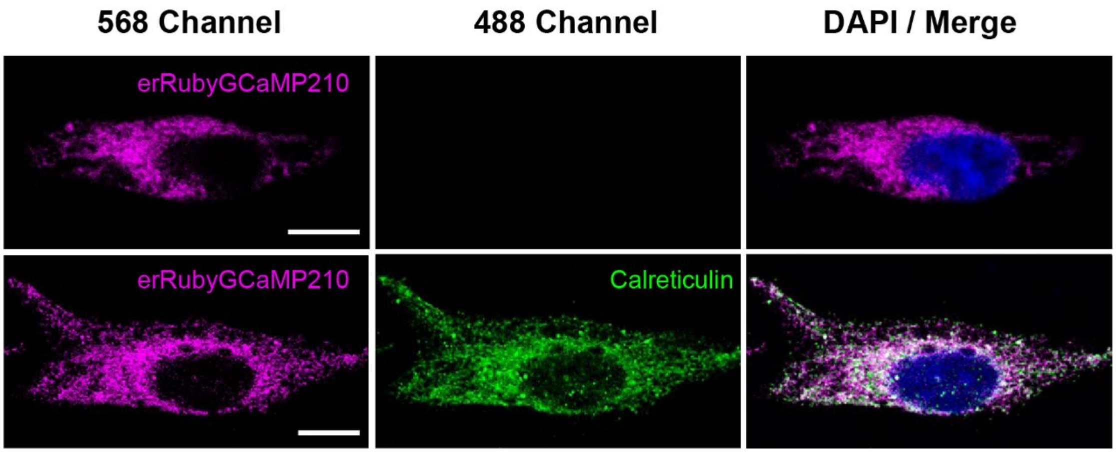


**Figure Supplement 2.**

**er-Ruby-GCaMP210 is a marker of the ER. (A)** Transiently transfected HeLa cells expressing er-Ruby-GCaMP210 display red fluorescence (magenta, 568 channel) and not green fluorescence (488 channel). **(B)** The pattern of cell expression of er-Ruby-GCaMP210 in the ER colocalized with that of calreticulin, an ER resident protein, detected with an anti-calreticulin antibody and revealed in green. Nuclei (in merge) were stained with DAPI. Scale bar for both images, 10 μm.

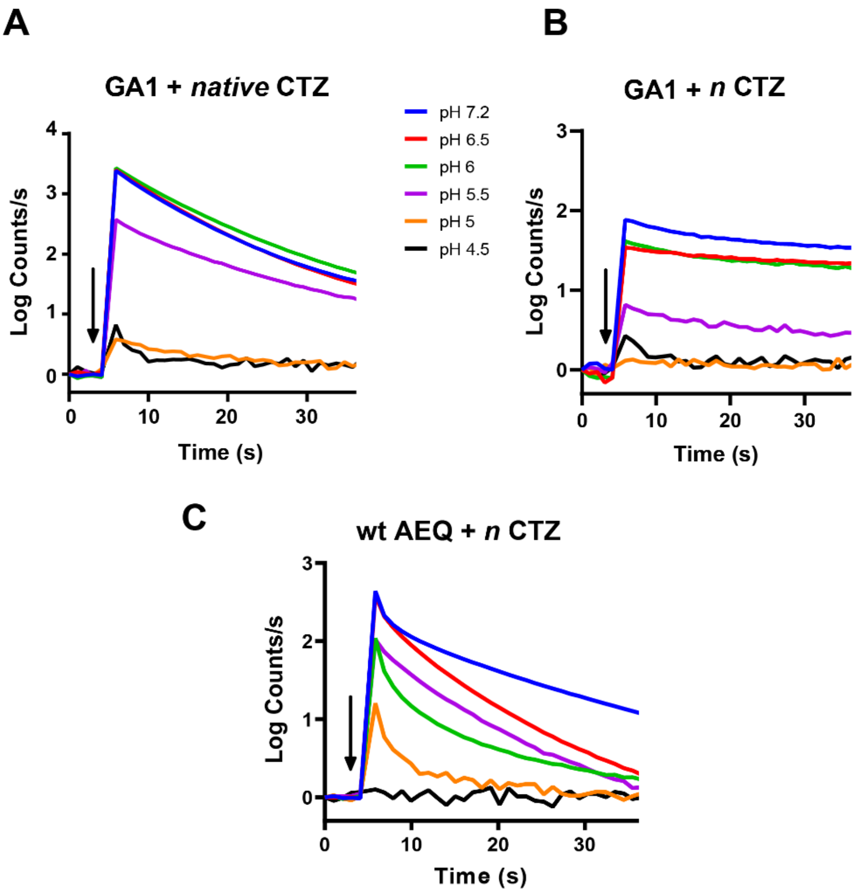


**Figure Supplement 3.**

**Effect of acidic pH on aequorin luminescence emission.** *E.coli* recombinant GA1 with either native- (**A**) or *n*-coelenterazine (CTZ) **(*B*)** at the indicated pHs for 20 min and then subjected to luminescence emission assay triggered with 10 mM $CaCl_2$ (arrow) and measured at the same pH. Buffers used were: acetate for range 4.5-5.5 pH; Na-MOPS for 6 and 6.5 pH; and Na-Hepes for 7.2 pH. Each trace is the value normalized with respect to its background emission before addition of calcium. Each trace is the average of 3-4 independent experiments. **(C)** *E.coli* recombinant wt-Aequorin reconstituted with *n*-coelenterazine.

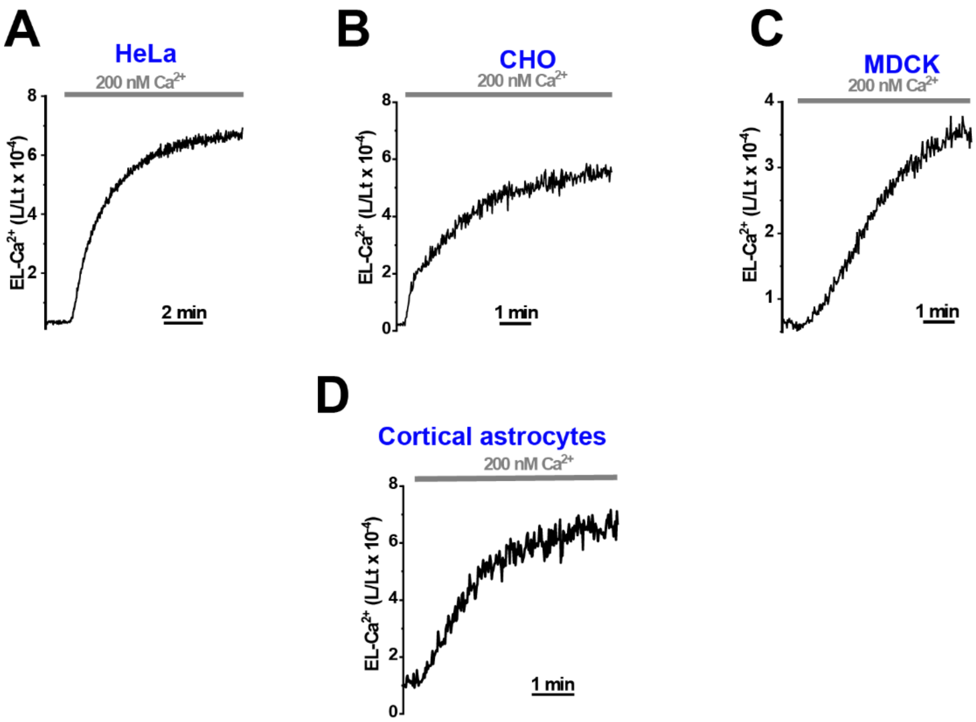


**Figure Supplement 4.**

**EL-$Ca^{2+}$ dynamics in various cell types.** HeLa, CHO, MDCK or cortical astrocytes were all transiently expressing ELGA1. For further details, refer to Figure 2.

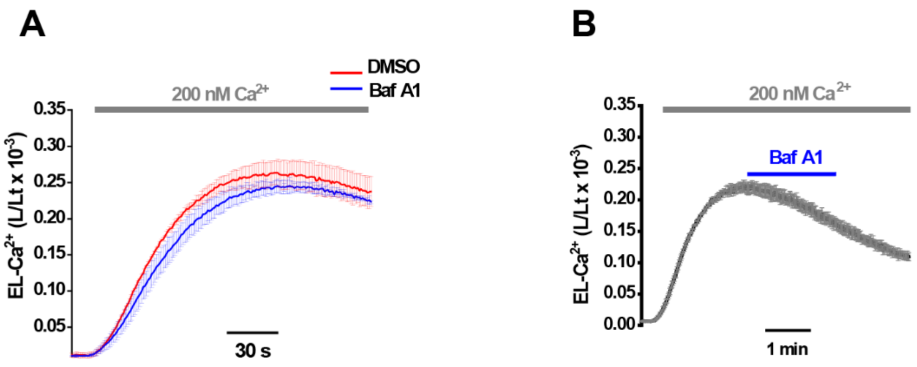


**Figure Supplement 5.**

**Effects of Bafilomycin A1 on EL-Ca$^{2+}$ dynamics. (A)** ELGA transiently expressing HeLa cells were preincubated with Bafilomycin A1 (1 μM, blue trace) or DMSO (red trace) for 1h. For further experimental details refer to Figure 2. **(B)** Acute application of Bafilomycin A1 (1 μM, 120s). Traces are the mean ± S.E.M. of 6 experiments from, at least, 2 independent days.

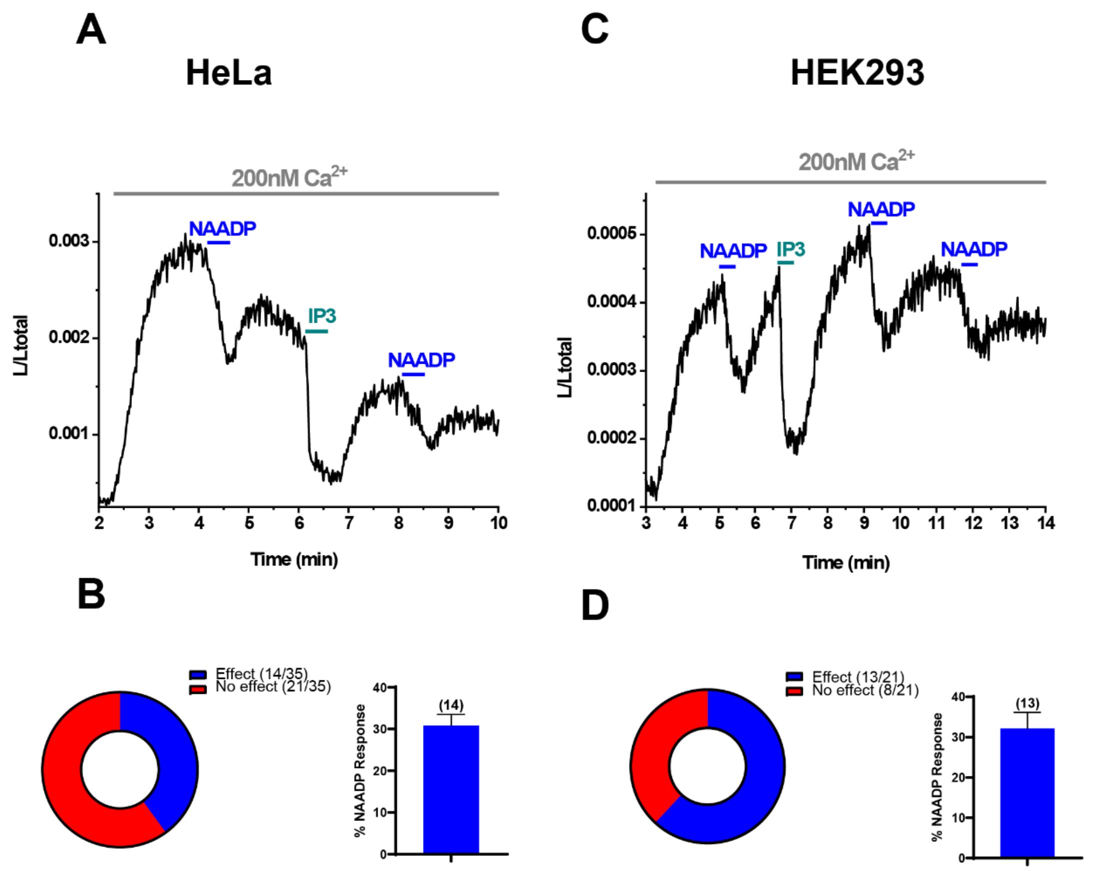


**Figure Supplement 6.**

**NAADP evoked $Ca^{2+}$ release from the endo-lysosome.** Representative measurements of stably-expressing ELGA in HeLa cells **(A)** or ELGA -transiently expressing HEK293T cells **(C)** in which NAADP (100 nM) provoked a response**.** $IP_3$ (2 µM) was also added for comparison. Statistical quantifications of the NAADP responses represented as percentages of total experiments performed and quantification of the NAADP release (% of EL-total $Ca^{2+}$ content at the steady-state) in the experiments in which a response was observed **(B, D).**

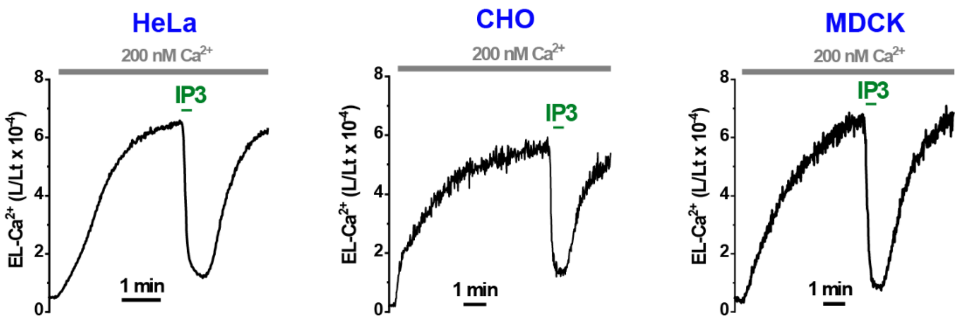


**Figure Supplement 7.**

**Responses to $IP_3$ in various cell types.** Representative responses to $IP_3$ (2 µM) in HeLa, CHO and MDCK cells expressing ELGA1. For further details, refer to Figure 3.

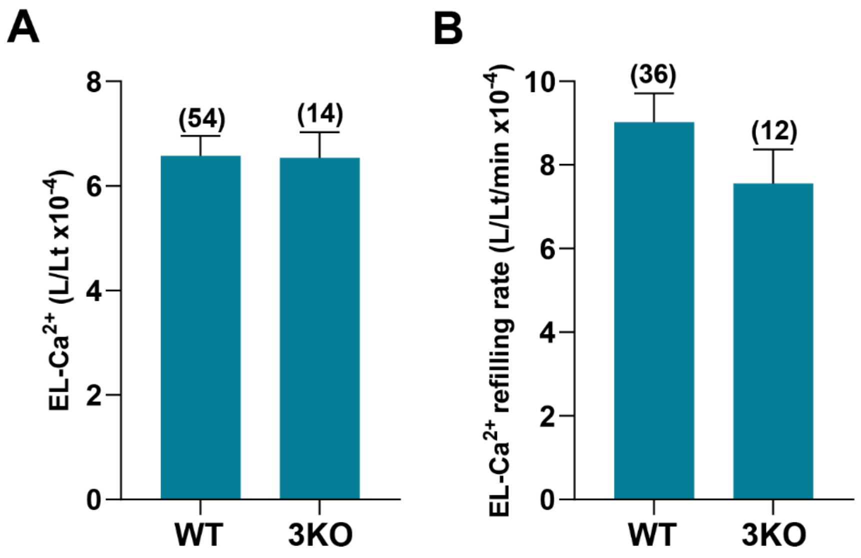


**Figure Supplement 8.**

**IP3R is not essential for EL-Ca$^{2+}$ refilling.** Comparison of resting $[Ca^{2}]_{EL}$ **(A)** and EL-refilling Ca$^{2+}$ rate **(B)** between control wt- and 3KO-HEK cells**.** A Student's *t* test was applied. Values are expressed as mean ± S.E.M. No significant differences were observed.

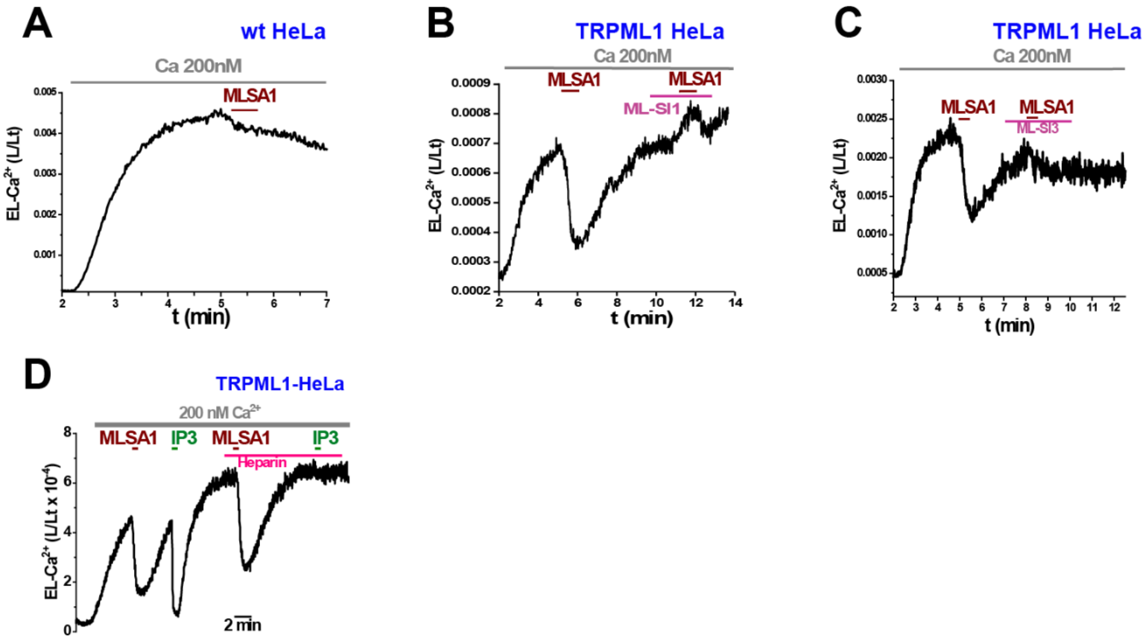


**Figure Supplement 9.**

**TRPML1 releases $Ca^{2+}$ from the EL-$Ca^{2+}$ store in HeLa cells. (A)** ELGA expressing HeLa cells were challenged with ML-SA1 (20 µM, 30 s). **(B, C)** Similar to protocol in (A), but in transiently expressing-TRPML1 HeLa cells. The second application of ML-SA1 was performed in the presence of ML-SI1 (10 µM) (B) or ML-SI3 (10 µM) (C). (**D)** Comparison between the ML-SA1 and the $IP_3$ responses.

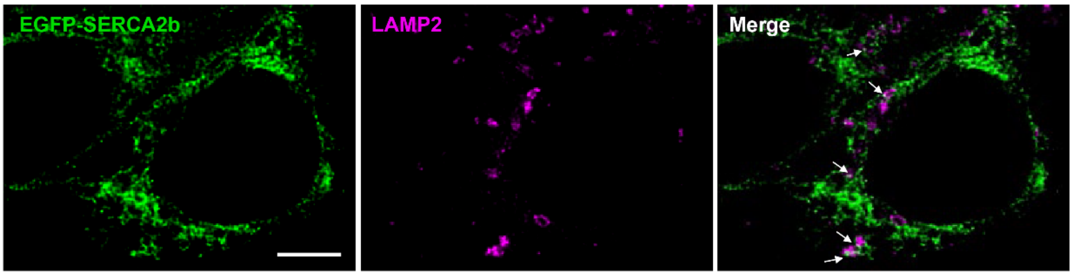


**Figure Supplement 10.**

**Part colocalization of SERCA2b with endo-lysosome marker.** Detail of one Z plane super-resolution image of HEK293T cells expressing EGFP-SERCA2b (green) with LAMP2 (magenta). Arrows indicate puncta of colocalization. Scale bar, 5 μm.

**Supplementary Movie 1.**

Confocal images, each corresponding to 0.38 μm, of MDCK cells transiently expressing ELGA.

**Supplement Table 1. Colocalization of ELGA with organelle markers.**

**Figure 1- supplement Table 1. Colocalization of ELGA with organelle markers.**

| **Organelle marker** | **Pearson coeficient (r)** | **Mander´s coefficient M1 (ELGA/marker)** | **Mander´s coefficient M2 (marker/ ELGA/)** | **N** |
|---|---|---|---|---|
| **LAMP-2** | 0.60 ± 0.03 | 0.55 ± 0.06 | 0.45 ± 0.04 | 10 |
| **ER** | 0.41 ± 0.03 | 0.36 ± 0.05 | 0.11 ± 0.02 | 14 |
| **Early endosome (EE)** | 0.27 ± 0.03 | 0.13 ± 0.01 | 0.24 ± 0.03 | 10 |
| **Mitochondria (mt)** | 0.20 ± 0.03 | 0.29 ± 0.05 | 0.14 ± 0.02 | 12 |

ELGA expressing HEK293T were fixed and: immunostained with the specific antibody anti-LAMP2; cotransfected with er-Ruby-GCaMP210 (ER); immunostained with the anti-early endosome antigen 1 (EE); or immunostained with the anti-TOM20 antibody (mt). Colocalization analysis of images acquired in a confocal microscopy. ***n*** indicates the number of cells analysed from 3 independent experiments.
